# High plasticity increases phenotype-environment mismatch leading to sub-optimal performance in underutilized crop species

**DOI:** 10.1101/2024.11.14.623330

**Authors:** Ganesh Alagarasan, Pieter A. Arnold, Eswarayya Ramireddy

## Abstract

As climate change threatens agricultural yields, crop diversification, including the reintroduction of underutilized species, has emerged as a proposed solution for resilience. But what determines the extent of environmental change that underutilized crop species can cope with upon their reintroduction? One critical factor is maladaptive plasticity, a theoretical concept suggesting that populations may respond negatively to rapid environmental shifts, at least in the short term. To test this empirically, we conducted a repeated-measures study using historically underutilized *Amaranthus* species as a model. Upon reintroduction, we observed higher plasticity in vegetative and reproductive traits than in life-history traits. This suggests that while these species adapt to immediate environmental changes through adjustments in growth and reproduction, limited plasticity in life-history traits may constrain long-term adaptation. Our study aimed to (1) identify the environmental factors driving these plastic responses, (2) determine whether these responses are maladaptive, and (3) assess if maladaptive plasticity also occurs within native environments. Using machine learning models, we found that temperature was a primary driver of plastic responses. Generalized Additive Model (GAM) reaction norm analysis further revealed these temperature-induced responses to be maladaptive, suggesting a mismatch between *Amaranthus*’ traits and the new environment. Applying transfer learning to predict responses in native settings, we found similar maladaptive responses to temperature, indicating that maladaptive plasticity may not be limited to non-native environments, thereby complicating climate adaptation efforts. This study underscore the need for detailed trait plasticity assessments before crop reintroduction for agricultural resilience.

## Introduction

Crop models predict significant yield reductions for major crops due to changing environmental conditions, particularly rising temperatures (Ray et al., 2015). For example, a one-degree Celsius increase in temperature can reduce wheat production by 6% and rice production by 10% (Peng et al., 2004; Asseng et al., 2015). To address these temperature-induced risks, crop diversification has emerged as a key climate change adaptation strategy, enhancing resilience and reducing the risk of yield losses (Gurr et al., 2016; He et al., 2023). As part of diversification, the reintroduction of underutilized crops, such as millet, is often posed as a sustainable solution for climate adaptation (Mayes et al., 2011; Li et al., 2020; Singh et al., 2022; Zonneveld et al., 2023). However, adapting to climate change presents unique challenges. Ecological principles suggest that species exposed to novel environmental conditions often face significant challenges in adapting, and this applies to cultivated crops as well (Chevin et al., 2010). For example, climate change is expected to expose crops in sub-Saharan Africa to such novel conditions, increasing production insecurity and increasing risks to food security in the region (Pironon et al., 2019). Thus, while underutilized crops offer potential resilience benefits, their reintroduction as a climate adaptation solution may sometimes be overestimated, as these crops also face significant adaptation barriers in new environmental contexts.

To ensure the successful reintroduction of underutilized crops as part of a diversification strategy, a mechanistic understanding of these crops’ responses to environmental stressors is essential. A key factor in this understanding is phenotypic plasticity—the ability of a crop to adjust its growth, development, and physiology in response to varying environmental conditions. Plasticity can be adaptive, where the response increases fitness, such as phosphorus starvation response (PSR) in rice adjusting root growth under phosphorus stress (Gamuyao et al., 2012). Alternatively, it can be maladaptive, where the response reduces fitness, as seen in soybean, where shading causes exaggerated stem elongation (ESE), leading to lodging and reduced yields in high-density planting (Qin et al., 2023). Theoretical studies suggest that species exposed to new environments may exhibit maladaptive plastic responses, where traits that were once beneficial may no longer align with the stresses posed by changing climates (Chevin and Lande, 2015; Chevin and Hoffmann, 2017). As a result, plasticity can lead to mismatches between genotype and environment, potentially hindering successful adaptation and long-term sustainability in novel ecological contexts. To empirically test this hypothesis, we selected *Amaranthus*, as many studies have previously highlighted that *Amaranthus* species have higher plasticity, although not thoroughly characterized (Zhang et al., 2020; Khanam and Oba, 2014; Chauhan and Abugho, 2012). Building upon this, we introduced *Amaranthus*, a historically underutilized crop species, into a new environment where it is not currently cultivated in contemporary agriculture.

The objectives of this study are to: (1) identify the environmental factors driving plastic responses in *Amaranthus*, (2) determine whether these responses are maladaptive, and (3) assess if maladaptive plasticity occurs solely in non-native environments or also in native ones. In field-level agricultural experiments, we use repeated-measures designs to capture trait-specific plasticity in response to reintroduction. To investigate the environmental drivers of trait-specific plastic responses in *Amaranthus*, we employed machine learning models. Specifically, we utilized Random Forest and Gradient Boosting models to identify and prioritize environmental factors that contribute to trait plasticity. To further characterize the nature of these plastic responses, we incorporated a GAM reaction norm analysis. This approach enabled us to map reaction norms—responses of specific traits to environmental gradients—and examine the functional form of plastic responses across various trait categories. To test whether plasticity observed in reintroduced populations also appears within native environments, we implemented transfer learning, a machine learning technique that leverages knowledge gained from one context to predict outcomes in another (Paymode and Malode, 2022; Zhao et al., 2022; Wang et al., 2023; Li et al., 2024). This approach allowed for a cross-environment comparison of trait plasticity, providing an empirical basis for assessing whether observed plasticity is an intrinsic characteristic of these populations or a context-dependent response. The novel methodology presented in this study—combining machine learning, GAM analysis, and transfer learning—offers potential applications across agriculture and conservation, from environment-based phenotype prediction and breeding program optimization to prioritizing species with maladaptive plasticity in biodiversity conservation. Underutilized crops are often celebrated as climate-resilient alternatives due to their adaptation to marginal conditions in native environments. However, our findings on maladaptive plasticity upon reintroduction challenge the assumption that these crops are inherently resilient and highlight the importance of thoroughly evaluating these crops’ adaptability beyond surface-level assumptions of resilience.

## Materials and methods

### (a) Plant materials and growth conditions

We obtained 88 genotypes of *Amaranthus* from the National Bureau of Plant Genetic Resources (NBPGR) in New Delhi (Table S1). While there is no concrete evidence to confirm their adaptation to high-altitude environments, *Amaranthus* are traditionally known to originate from and thrive in mountainous regions like the Himalayas and the Andes (Joshi, 1991; Blair et al., 2023; Sahagun, 1963; Tucker, 1986). Upon receipt, seeds from each genotype underwent germination testing to assess their viability. This was achieved by planting the seeds in small pots filled with coir pith, a suitable germination medium. The germination efficiency was carefully monitored to ensure the quality of the seed material. Following the germination assessment, the seeds were stored at a controlled temperature of 4°C to maintain their viability and vigour. Open field experiments were conducted at two locations: one situated at 13.7414°N, 79.6011°E (in Andhra Pradesh, India, altitude approx. 110 m) and the other at 12.1966°N, 78.3738°E (in Tamil Nadu, India, altitude approx. 420 m), from 2019 to 2021. The field was initially prepared for cultivation through ploughing, using a rotavator to effectively remove weed growth. After a week, ridges and furrows were created in the field. Each specific genotype’s seeds were then placed on top of these ridges and subsequently covered with soil. Throughout the growth cycle, the plants were regularly watered to ensure adequate moisture levels. To prevent any potential crop damage caused by pest attacks, insecticides were applied within two weeks of seed germination. To prevent overcrowding, which can impede plant growth and development, seedlings were thinned during the early stages of growth. Additionally, during the crop growth period, any admixture that may have occurred was carefully removed. It is important to note that each genotype consisted of approximately 150 to 200 plants until reaching maturity. This allowed for a sufficient sample size to assess and evaluate the performance of each genotype throughout the entire growth cycle. Plant height and stem girth measurements were taken before the onset of flowering. The flowering date was determined when at least half of the plants within a plot had begun flowering. Following the cessation of inflorescence growth, the length and number of branches in the inflorescence were measured. The maturity days were calculated by recording the number of days from seed sowing to the harvest day. All measurements were conducted manually.

### (b) Plasticity estimation

The genotypic plasticity score (hereinafter referred to as ‘GenoPlast score’) was calculated to assess the phenotypic plasticity of each genotype, defined mathematically as 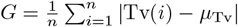, where Tv(*i*) represents the trait value for a particular condition *i, µ*_Tv_ denotes the median trait value across all conditions, |Tv(*i*) − *µ*_Tv_|is the absolute difference between the trait value for condition *i* and the median trait value (ensuring that all differences are treated as positive values), and *n* refers to the total number of conditions.

### (b) Machine learning models

To identify and prioritize the environmental factors influencing phenotypic plasticity in *Amaranthus* genotypes, we applied machine learning techniques, specifically Random Forest (RF) and Gradient Boosting (GB), to model the relationship between climatic conditions and plant traits. The dataset used for model training was derived from field experiments, where climatic variables (e.g., maximum and minimum temperature, rainfall, wind speed, humidity, UV index, and sunny days) were treated as input features, and phenotypic traits (e.g., plant height, stem girth, inflorescence length, number of branches, flowering time, and time to maturity) were the target variables. The climatic data during the experimental period (Location 1: Jun to Sep 2019 and Jan to Apr 2021; Location 2: Jun to Sep 2020 and Jan to Apr 2021) was collected from https://www.worldweatheronline.com/. Both models were trained after applying a `StandardScaler` (a function of sklearn library in python) to ensure uniform scaling of the features, which is particularly important for models like Gradient Boosting. The Random Forest model, a bagging ensemble method, aggregates predictions from multiple decision trees, each trained on a bootstrapped sample of the data. The final prediction is made by averaging the outputs of all trees when performing regression. Mathematically, this can be expressed as 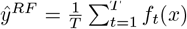, where *ŷ*^*RF*^ is the predicted value, *T* is the number of trees, and *f*_*t*_(*x*) is the prediction from the *t*-th tree for input *x*. The Gradient Boosting model, on the other hand, builds trees sequentially. Each tree tries to minimize the residual error from the previous trees by fitting the model to the residuals, which are calculated as the negative gradient of the loss function. This process can be represented as 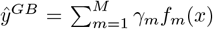, where *ŷ*^*GB*^ is the predicted value, *M* is the number of boosting iterations, and *γ*_*m*_ is the weight assigned to the *m*-th tree. After training, feature importances were calculated for each model to assess the relative contribution of each climatic variable in predicting the phenotypic traits. These importance scores were averaged across all traits to identify the most influential climatic factors.

### (c) Reaction norm analysis

For each trait *Y* (e.g., plant height, stem girth, flowering time), the model was specified as *Y*_*i*_ = *β*_0_ + *s*(Climatic Factor_*i*_) + Genotype_*i*_ + *ϵ*_*i*_, where *Y*_*i*_ represents the phenotypic trait for the *i*-th observation, *β*_0_ is the intercept term, *s*(Climatic Factor_*i*_) is the smooth term for the most influential climatic factor, modeled using penalized splines to capture its non-linear effect, Genotype_*i*_ is a factor representing the genotype, and *ϵ*_*i*_ is the error term. The smooth function *s*(Climatic Factor_*i*_) allows the relationship between the climatic factor and the trait to be non-linear, providing flexibility to model complex patterns in the data. The model was fitted using the fREML method (Fast Restricted Maximum Likelihood), which estimates the smoothness of the splines and allows for efficient computation. Predictions were made for both population-level and genotype-specific responses, with the results transformed back to the original scale of the data after centering and scaling for numerical stability. Model performance was assessed through residual analysis and diagnostic plots, including normality checks of the residuals. We also reported key model fit statistics such as AIC (Akaike Information Criterion), BIC (Bayesian Information Criterion), and explained deviance, which provide a measure of how well the model fits the data.

### (d) Transfer learning approach

We incorporated a transfer learning approach to enable our GAM model to generalize across both internal (experimental) and external datasets. The external dataset, derived from a study by Raiger et al. (2014), included phenotypic data on plant height for 54 Amaranth genotypes in their native habitat, providing environmental variability absent in the internal dataset. This transfer learning setup aimed to enhance the model’s adaptability to new, unobserved environmental conditions. For transfer learning, we combined both datasets into a single input, *D* = *D*_int_ ∪ *D*_ext_, where 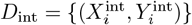 represents the internal dataset, and 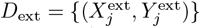 represents the external dataset. Here, *X* includes predictors (e.g., climatic factors, genotype), and *Y* represents the response variable (e.g., plant height). We aligned feature scales and distributions through centering and scaling variables like growth temperature and plant height to ensure consistency across *D*_int_ and *D*_ext_. The model was then specified similarly to the initial GAM, modeling phenotypic traits as functions of influential climatic factors (e.g., temperature) and genotype. This combined model enabled us to predict both population-level and genotype-specific responses across datasets, leveraging shared patterns while accommodating environmental differences. Model performance was assessed through residual analysis, normality checks, and summary statistics, including AIC, BIC, and explained deviance.

## Results

### (a) Differential trait plasticity

To identify key traits influencing plasticity and discern common patterns across traits, as well as to determine which genotypes exhibit similar responses to environmental conditions, we employed the k-means algorithm, an unsupervised machine learning technique, and principal component analysis (PCA). Using phenotypic data obtained under various experimental conditions, we quantified the GenoPlast score. The genotypes were grouped into distinct clusters using the k-means algorithm, where optimal clustering was confirmed by the maximum average silhouette score, indicating the formation of four distinct clusters (Figure 1A). PCA of these clusters revealed common patterns of plasticity across traits, with ‘Plant height’ (36.21%) and ‘Stem girth’ (23.94%) being the most significant contributors to the overall variance. ‘Flowering’, despite contributing to only 4.67% of the total variance, exhibited the strongest loading on PC2, underscoring its specific importance in explaining variance along this component. The four primary traits — ‘Plant height’, ‘Stem girth’, ‘Inflorescence length’, and ‘Inflorescence branch number’ — collectively accounted for 94% of the total variance, underscoring their potential as key differentiators among genotypes (Figure 1B). The distinct clusters, identified through k-means, show that common patterns of plasticity are shared among certain genotypes in principal component space (Figure 1B), where the clustering may reflect unique evolutionary or cultivation histories. The cluster structures offer a more precise map of the traits and genotypes that are most responsive to environmental changes, emphasizing their importance in the ecological adaptability of *Amaranthus* (Figure 1C). The plasticity patterns observed in genotypes offer guidance for targeted breeding and conservation efforts. For example, genotypes from cluster 2 may be prioritized for adaptability in flowering times, while clusters 1, 3, or 4 may be considered for versatility in both flowering and maturity (Figure 1C). Next we examined the plasticity of individual traits across the entire set of genotypes, regardless of their cluster classification. This approach aimed to identify the trait(s) demonstrating the highest degree of plasticity, which would indicate the primary plastic trait within the population. We found that inflorescence length and plant height exhibited the highest degree of plasticity (Figure 1D). Conversely, the number of days to reach maturity showed minimal or no plasticity, suggesting it may be more strongly determined by non-plastic genetic factors (Figure 1D). Following the analysis of individual traits, we calculated the composite GenoPlast score across all six traits for each genotype, finding the most plastic genotypes within the population (Figure 1E). These findings not only highlight the significant plastic traits in *Amaranthus* genotypes but also underscore the importance of targeted trait selection for adaptability in varying climates. But what are the key environmental variables driving these plastic responses? And how can machine learning models help us identify them? In the following section, we uncover these insights through modeling techniques.

**Figure 1:**
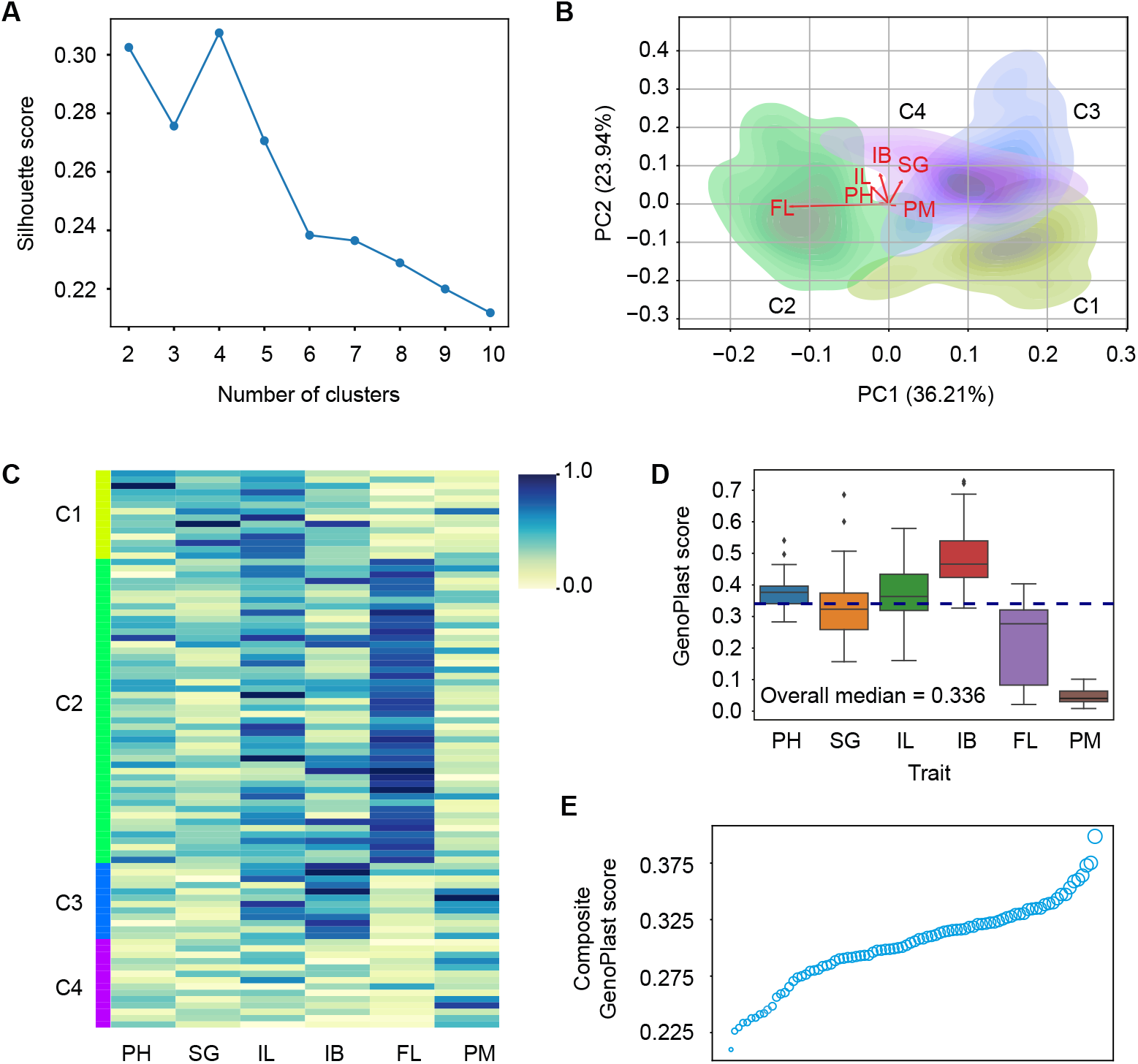
Genotype clustering across *Amaranthus* spp. and their trait plasticity. (A) A silhouette plot displaying average silhouette scores for various cluster numbers, identifying the optimal cluster number (four) by the highest score. (B) Biplot of first two principal components (PC1 and PC2) and trait vectors from Principal Components Analysis (PCA) of six phenotypic traits (I.B = Inflorescence Branch Numbers; PH = Plant Height; IL = Inflorescence Length; SG = Stem Girth; FD = Flowering Date ; PM = Plant Maturity) including density contours by each of the four clusters that show genotypes’ distribution in principal component space. (C) Combined heatmaps for each cluster, with colour intensity indicating GenoPlast score. (D) Box plots representing the GenoPlast score for each phenotypic trait across all genotypes. The dashed blue line marks the median value across all traits.(E) A bubble plot depicting genotypes’ composite GenoPlast scores, where bubble size corresponds to score value.

### (b) Climatic variables triggering plasticity

To evaluate and prioritize the environmental factors influencing phenotypic plasticity in *Amaranthus* genotypes, we trained Random Forest and Gradient Boosting models using climatic data as input features and plant traits as target variables. The feature importances derived from these models were averaged across all target traits to determine the relative influence of each climatic variable. The analysis revealed that in the RF model, the most significant climatic variables were Max temp (0.13), Avg temp (0.13), and Cloud % (0.11). In comparison, the GB model identified Max temp (0.14), Cloud % (0.14), and Humidity % (0.13) as the top features (Fig. 2). Both models consistently highlighted Max temp and Avg temp as critical variables across multiple traits, demonstrating the substantial impact of temperature on phenotypic responses. Notably, the GB model placed greater emphasis on Max wind (0.13) and Sun days (0.13) than the RF model, suggesting that these factors may contribute more nuanced influences when models are built sequentially (Fig. 2). To gain deeper insights, we also examined the impact of each climatic variable on individual traits (Fig. S1). This detailed analysis revealed that Max temperature was a leading predictor across traits like plant height, stem girth, inflorescence length, and time to flowering, underscoring its broad influence on plant growth and development. Cloud cover and average temperature were prominent for traits such as height, number of branches, and stem girth, suggesting that both overall temperature and light availability are key drivers of plasticity in *Amaranthus* genotypes. Notably, the GB model placed greater emphasis on Max wind and Sun days for traits like stem girth and flowering time, suggesting these factors contribute nuanced influences better captured by sequential learning models (Fig. S1). This systematic calculation underscored that Max temp, Avg temp, and Cloud % were consistently influential in driving plasticity, highlighting their pivotal role in the environmental responsiveness of *Amaranthus* genotypes. However, whether the induced plasticity is adaptive or maladaptive in response to temperature gradients remains an open question. To address this, we turn to GAM model in the following section to provide insights into the potential adaptiveness of these plastic responses.

**Figure 2:**
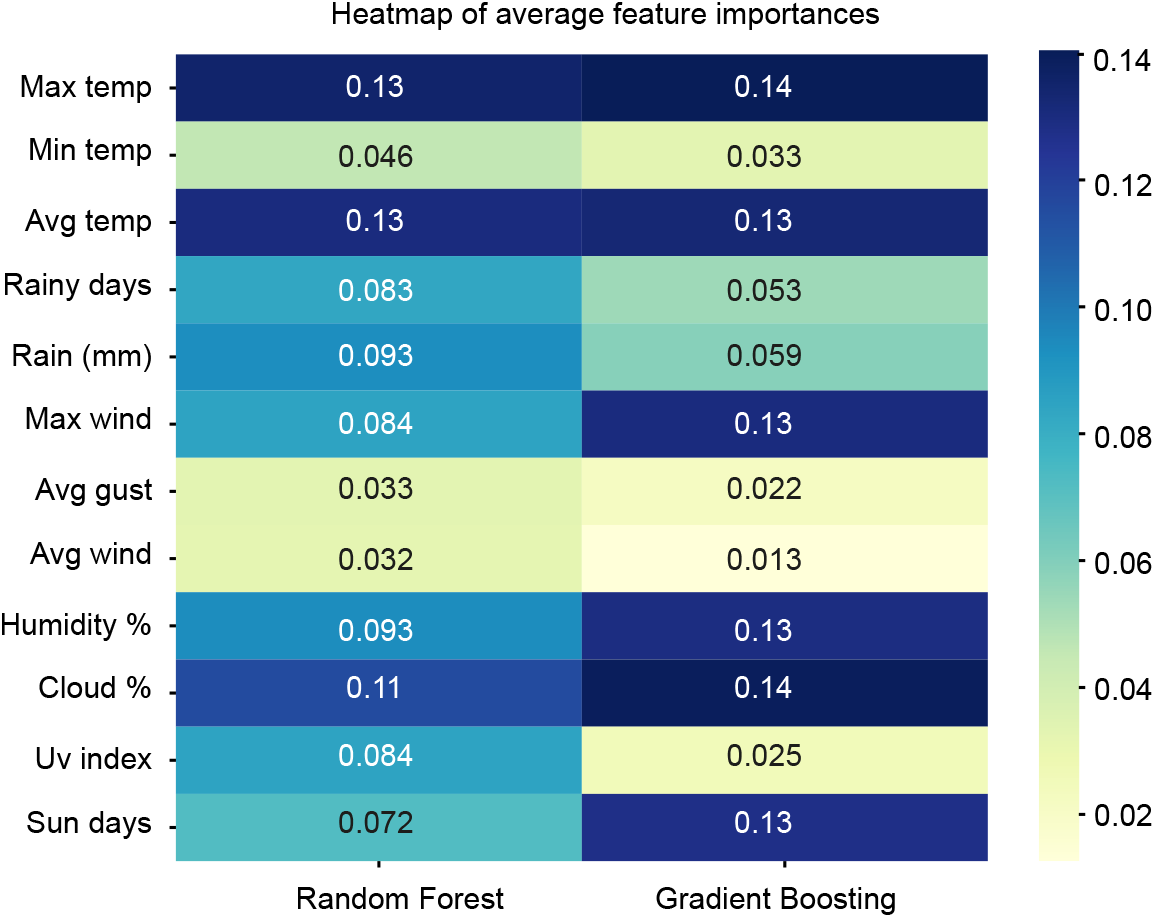
Climatic variable importance hierarchy revealed by Random Forest and Gradient Boosting models for plant trait prediction. This heatmap illustrates the average feature importances derived from Random Forest and Gradient Boosting models for predicting various plant traits. The climatic variables, encompassing aspects like temperature, precipitation, wind, humidity, cloud cover, UV index, and sunshine duration, were employed as predictive features. The color gradient, ranging from warmer to cooler hues, signifies the relative significance of these climatic variables in predicting plant traits.

### (c) Insights from reaction norm model

The GAM analysis revealed negative correlations between vegetative traits and temperature. Specifically, the model demonstrated an exceptionally high R2 value of 0.958 for plant height, indicating that the model captures the significant influence of both temperature and genotype (Fig. 3A; residual and Q-Q plots in Fig. S2)(Table S2). The analysis identified a non-linear relationship, with an optimal temperature range facilitating maximal plant height, beyond which further temperature increases could detrimentally impact growth, suggesting a threshold at which temperature triggered plasticity becomes maladaptive for the plant. The model fit for stem girth was similarly robust, with an R2 of 0.902 (Fig. 3B). Statistically significant effects from temperature (*P <* 0.001) suggest a non-linear response, indicating a corresponding optimal temperature range that supports stem girth development. The consistency of these patterns in both vegetative traits—plant height and stem girth—highlights the effect of temperature on the structural development and vigour of the plant that underpins growth dynamics with environmental change. These findings emphasize the potential for thermal stress to induce maladaptive plasticity, which could compromise growth processes, biomass accumulation, and plant structural integrity that are key components of plant fitness under agricultural settings. In examining the reproductive traits, the GAM analysis for inflorescence branch number (Fig. 4A) and inflorescence length (Fig. 4B) revealed a negative effect of temperature on these traits. For inflorescence branch number, the model had a good fit (R2 = 0.811) but was slightly less predictive compared to the two vegetative traits (Fig. S2). Inflorescence length was significantly influenced by temperature (*P <* 0.001), indicating that it is affected by temperature with a non-linear relationship. Inflorescence branch number showed temperature as highly significant factor (*P <* 0.001), and sound model fit (R2 = 0.784). The relationship between temperature and the number of inflorescence branches was also non-linear, suggesting an optimal temperature range for maximizing branch number.

**Figure 3:**
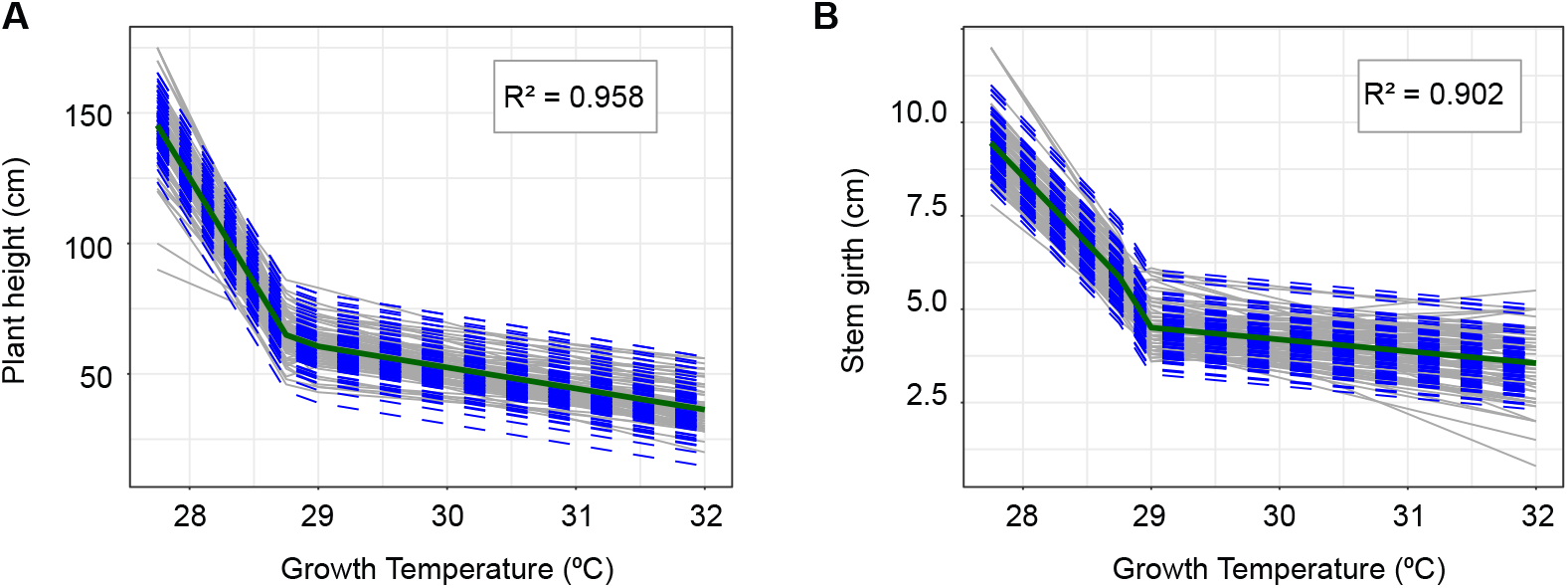
Vegetative trait plasticity. (A) Plant height and (B) stem girth are plotted as a function of growth temperature. The trait values are plotted (grey lines) along with predicted trait values derived from generalized additive model (GAM) in non-native environments. Dashed blue lines represent genotype-specific predictions, while a solid dark green line shows population-level predictions.

**Figure 4:**
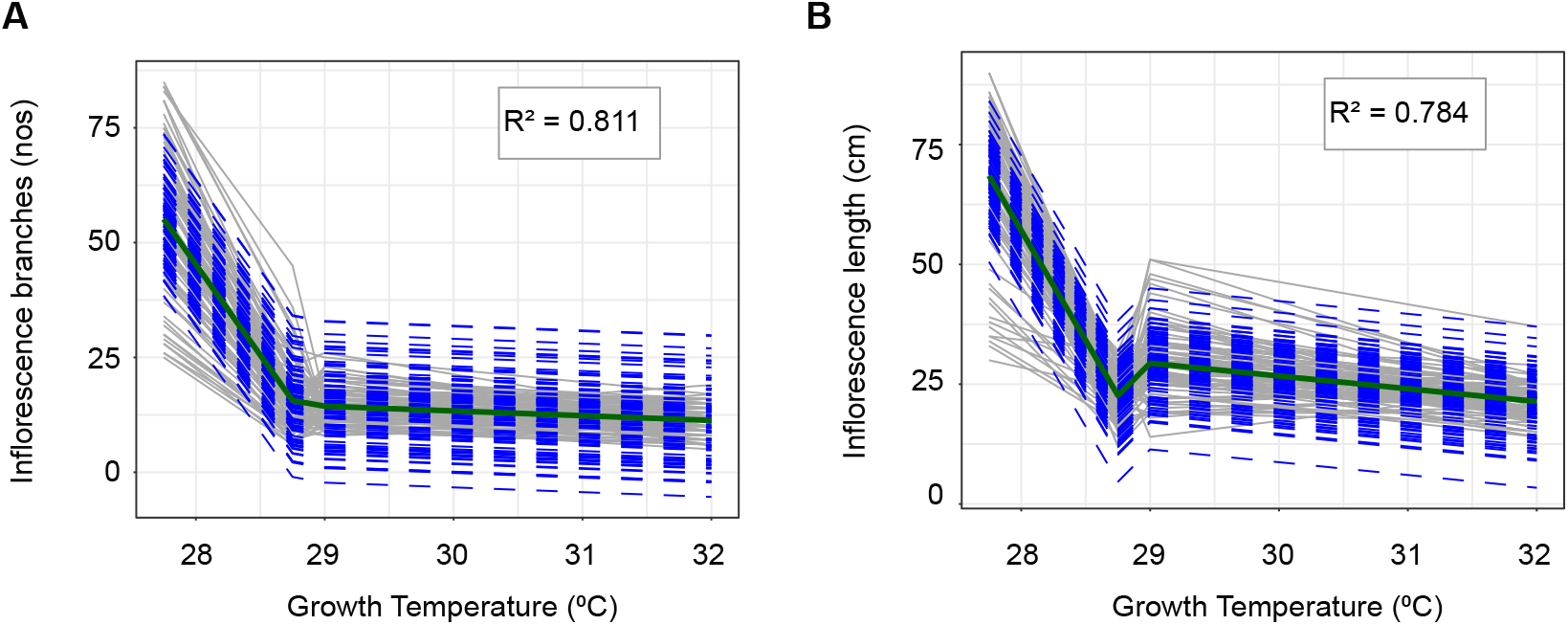
Reproductive trait plasticity. A) number of inflorescence branches, (B) inflorescence length, are plotted as a function of growth temperature. The trait values are plotted (grey lines) along with predicted trait values derived from generalized additive model (GAM) in non-native environments. Dashed blue lines represent genotype-specific predictions, while a solid dark green line shows population-level predictions.

The timing of flowering and maturity were also affected by temperature. For the flowering trait model (R2 = 0.778), significant smooth term for temperature (*P <* 0.001) indicates that this factor have a crucial role in determining the timing of flowering (Fig. 5A; Fig. S2). Despite the absence of a clear non-linear relationship that was found in vegetative and reproductive traits, flowering demonstrates a certain degree of variability among genotypes, with some showing extremely early flowering in response to thermal stress. This suggests that while temperature is a significant predictor of flowering time, the response is not uniform across genotypes. The model for time to maturity (R2 = 0.796) had significant smooth term for temperature (*P <* 0.001) similar to time to flowering (Fig. 5B). The consistent pattern of trait value decline across increasing temperatures points to maladaptive plasticity in the genotypes of *Amaranthus* species that we studied. Rather than providing a means to cope with and survive in warmer conditions, higher plasticity actually compromised the plants’ structural integrity, growth, and reproductive success. These findings underscore the potential vulnerability of underutilized crops to climate change, particularly to rising temperatures, and highlight the importance of considering both the direction and magnitude of plastic responses in future resilience and adaptation strategies. This raises an important question: Is maladaptive plasticity in response to increasing temperatures specific to non-native environments, or is it also observable in native environments? In the following section, we provide insights using transfer learning modeling techniques to address these questions.

**Figure 5:**
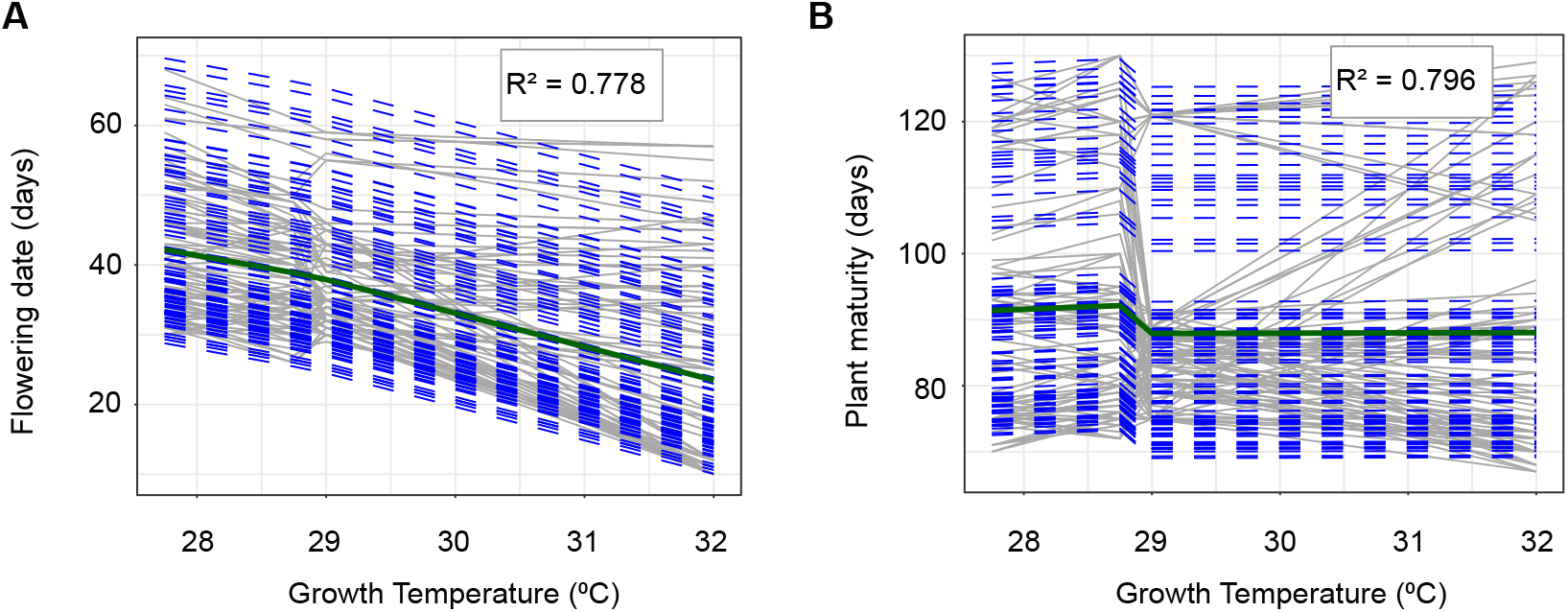
Life-history trait plasticity. (A) flowering days, (B) plant maturity, are plotted as a function of growth temperature. The trait values are plotted (grey lines) along with predicted trait values derived from generalized additive model (GAM) in non-native environments. Dashed blue lines represent genotype-specific predictions, while a solid dark green line shows population-level predictions.

### (d) Insights from transfer learning-based GAM model

The transfer learning GAM model demonstrated strong predictive capability for plant height in the native environment, achieving an adjusted *R*^2^ value of 0.742 (Fig. 6A and 6B). The QQ plot and smooth term plot support the model’s validity and distribution assumptions (Fig. S3).The model also explained 82.2% of the total deviance, highlighting its robust ability to capture the complexity of plant height responses across different environmental conditions. The model fit statistics included an fREML score of 493.21 and a scale estimate of 1.2373. Additionally, the AIC and BIC values of 1387.543 and 1980.359, respectively, support the model’s adequacy by balancing goodness of fit with model complexity. A significant contrast was observed in plant height responses between native and non-native environments (Fig. 6B). Specifically, plants grown in the native environment maintained consistently reached taller heights, even under conditions of elevated temperatures. The minimum height observed in the native environment surpassed the maximum height recorded in the non-native setting (Fig. 3A and 6A). This suggests that plants possess a distinct growth advantage or resilience when in their native habitats. This finding underscores that these maladaptive traits are intrinsic to the genotypes themselves, rather than being influenced by unique ecological conditions or the structure of the experimental setup.

**Figure 6:**
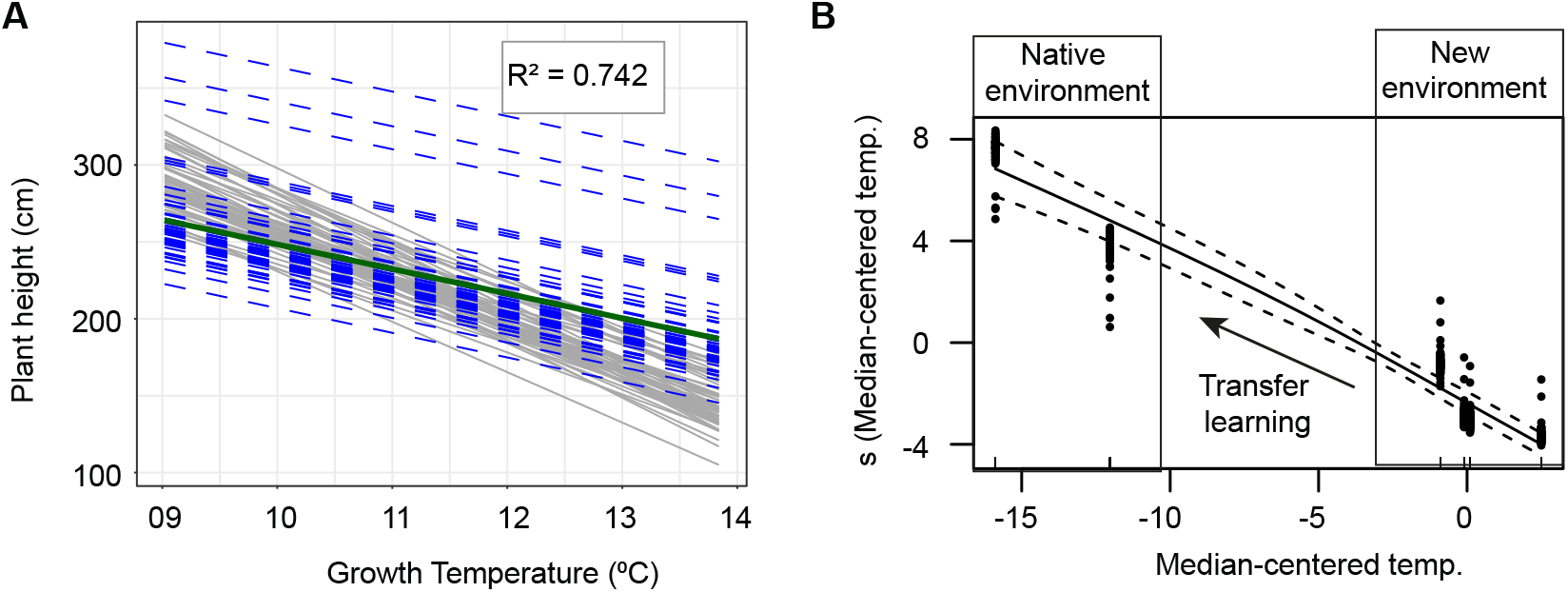
Transfer learning-based prediction of plant height plasticity in amaranth genotypes. The experimental data from non-native environments were used to train the model, which predicts the plasticity of various genotypes in their native environments. Plant height plasticity (A) is plotted as a function growth temperature, with residual plots (B). The trait values are plotted (grey lines) along with predicted trait values derived from generalized additive model (GAM). Dashed blue lines represent genotype-specific predictions, while a solid dark green line shows population-level predictions.

## Discussions

Our study aimed to evaluate the potential of reintroducing historically underutilized crops, using *Amaranthus* as a model organism. Specifically, we focused on a less-explored aspect in crop adaptation studies: phenotypic plasticity. We examined whether phenotypic plasticity aids or hinders adaptation during crop reintroduction. To achieve this, we employed machine learning models and GAM model for a detailed analysis of the functional aspects of plasticity. While the common understanding in crop science is that plasticity enables crops to buffer against environmental changes, the functional implications of this trait remain underexplored. Interestingly, our study reveals that plasticity is not always adaptive. Excessive plasticity can lead to a mismatch between phenotype and environment, resulting in suboptimal performance and challenging successful crop reintroduction.

The plasticity score proposed in our study, termed the GenoPlast score, is straightforward and effective for comparing phenotypic plasticity across multiple traits and species. This metric revealed substantial genetic variation in plasticity among the *Amaranthus* genotypes (Fig. 1). Like other analytical methods, the GenoPlast score can be applied to prioritize genotypes in breeding programs based on their trait-specific plasticity. For instance, it can help identify genotypes that exhibit high plasticity in desirable traits, while maintaining minimal or less plasticity in other traits. This targeted approach supports the selection of genotypes that balance adaptability with stability, optimizing their performance for specific environmental challenges.

One of the common challenges in field experiments is the multitude of factors influencing trait responses. While trait variability is often reported, the underlying reasons for this variability receive less attention. Employing machine learning techniques can be particularly valuable in such cases, as they help identify the primary factors driving variability in specific traits across different field trials. For example, in this study, the use of Random Forest and Gradient Boosting models allowed us to identify key drivers of plasticity (Fig. 2). These models highlighted the significant impact of temperature and other environmental variables, providing a clearer understanding of the factors influencing phenotypic plasticity.

While the general responses of crops to heat stress are well-documented, the functional form of plasticity—the way plants modulate their growth and development in response to varying temperatures—remains less defined. Functional plasticity involves understanding not just the existence of trait variability, but how the magnitude and direction of these responses change under specific environmental conditions. GAMs offer a powerful tool for quantifying this functional form, as they can capture non-linear relationships between environmental variables and phenotypic traits. GAMs can effectively measure the extent of plasticity (its magnitude) and indicate whether this plastic response is adaptive or maladaptive (its direction). Additionally, GAMs can predict phenotypic outcomes based on environmental cues, providing insights into potential performance under future climate scenarios. In our study, we employed GAMs and tranfer learning derived GAM model to reveal how traits such as vegetative and reproductive traits respond to temperature gradients (Fig. 3, 4, 5 and 6). This approach not only allowed us to quantify plasticity in these traits but also helped identify the temperature thresholds beyond which plastic responses turned maladaptive. The insights gained from this analysis highlighted that excessive plasticity, driven by high temperatures, could result in a phenotype-environment mismatch, ultimately challenging successful crop reintroduction. Given this context, it becomes clear why the expansion of underutilized crops like quinoa is significantly constrained by higher temperatures (Bazile et al., 2016; Tovar et al., 2020). To address this limitation, various studies have been exploring the genetic basis for heat tolerance (Tovar et al., 2020; Xie et al., 2023; Zhang et al., 2024). However, as our study highlights, if the majority of traits in these crops exhibit high plasticity, then it is essential to place greater emphasis on exploring the nature and implications of this plasticity. Determining whether the observed plasticity is adaptive or maladaptive is essential for guiding breeding programs and developing strategies to enhance resilience to temperature stress, aligning with the evolutionary principles discussed here (Romero-Mujalli et al., 2021; Acasuso-Rivero et al., 2019; Fox et al., 2019; Brooker et al., 2022).

In field experiments, extreme climatic conditions often result in complete crop failure, leading to insufficient datasets. In our field experiments conducted between 2019 and 2021, we observed this firsthand, experiencing complete crop failures more than six times due to the high plasticity of *Amaranthus* species under severe rainfall and heat stress. Transfer learning is particularly useful in such cases. For example, in our study, we used transfer learning to analyze plant height in *Amaranthus* genotypes in their native environment, where data availability was limited. This analysis provided an important insight: maladaptive plasticity is not a result of experimental conditions but rather an inherent trait in *Amaranthus* (Fig. 6). Overall, our analysis demonstrates that the adaptive behaviours of underutilized crops often embody the Red Queen’s hypothesis of evolutionary catch-up (Van Valen, 1973). This evolutionary theory proposes that organisms must continuously adapt and evolve to gain an advantage and keep pace with the changing environment. This continuous evolutionary process is crucial for survival, as static or maladaptive traits can lead to a decline in fitness over time. In our study on orphan crops, the observed phenomenon of maladaptive plasticity underlines this principle. It suggests that these crops, despite their evolutionary history of adaptation to niche environments, now face new challenges that may outpace their ability to adapt under current rates of environmental change. In this context, understanding reaction norms and the evolution of plasticity is vital, considering the new selection pressures and tradeoffs they introduce in plant traits, as discussed in depth by Van Kleunen and Fischer (2005), Matesanz et al. (2010), Chevin et al. (2010), Ghalambor et al. (2007), Murren et al. (2015), and Schlichting (1986). In conclusion, our results imply that, despite the initial promise of using underutilized crops like *Amaranthus* for climate adaptation, the potential for maladaptive plasticity must be considered, as it could hinder resilience under fluctuating environmental conditions. The analytical framework presented in this study, which incorporates machine learning models for identifying crucial factors controlling plasticity, GAM model reaction norm analysis, and transfer learning, opens new avenues for exploring functional form plasticity in crop adaptation research.

## Acknowledgements

We are grateful to the numerous individuals who have contributed to the field work. Special thanks to Pavan Kumar Avilala for assisting with extensive field trials. We also acknowledge the Research Advisory Committee for G.A., at the Indian Institute of Science Education and Research – Tirupati, India, for their conceptual discussions. We thank Prof. R.P. Sharma and Prof. Subha Srinivasan for commenting on the initial draft. This work received support from the Department of Biotechnology, India, under Grant No. BT/PR23613/BAP/118/354/2017, awarded to E.R. A.G. is the recipient of an IISER Tirupati Institutional PhD Fellowship.

## Author contributions

Ganesh Alagarasan: Conceptualization; data curation; formal analysis; investigation; methodology; software; validation; visualization; writing – original draft; writing – review and editing. Pieter A. Arnold: Methodology; validation; visualization; writing – review and editing. Eswarayya Ramireddy: Conceptualization; funding acquisition; investigation; methodology; project administration; resources; supervision; writing – original draft; writing – review and editing.

---

**Fig. S1:**
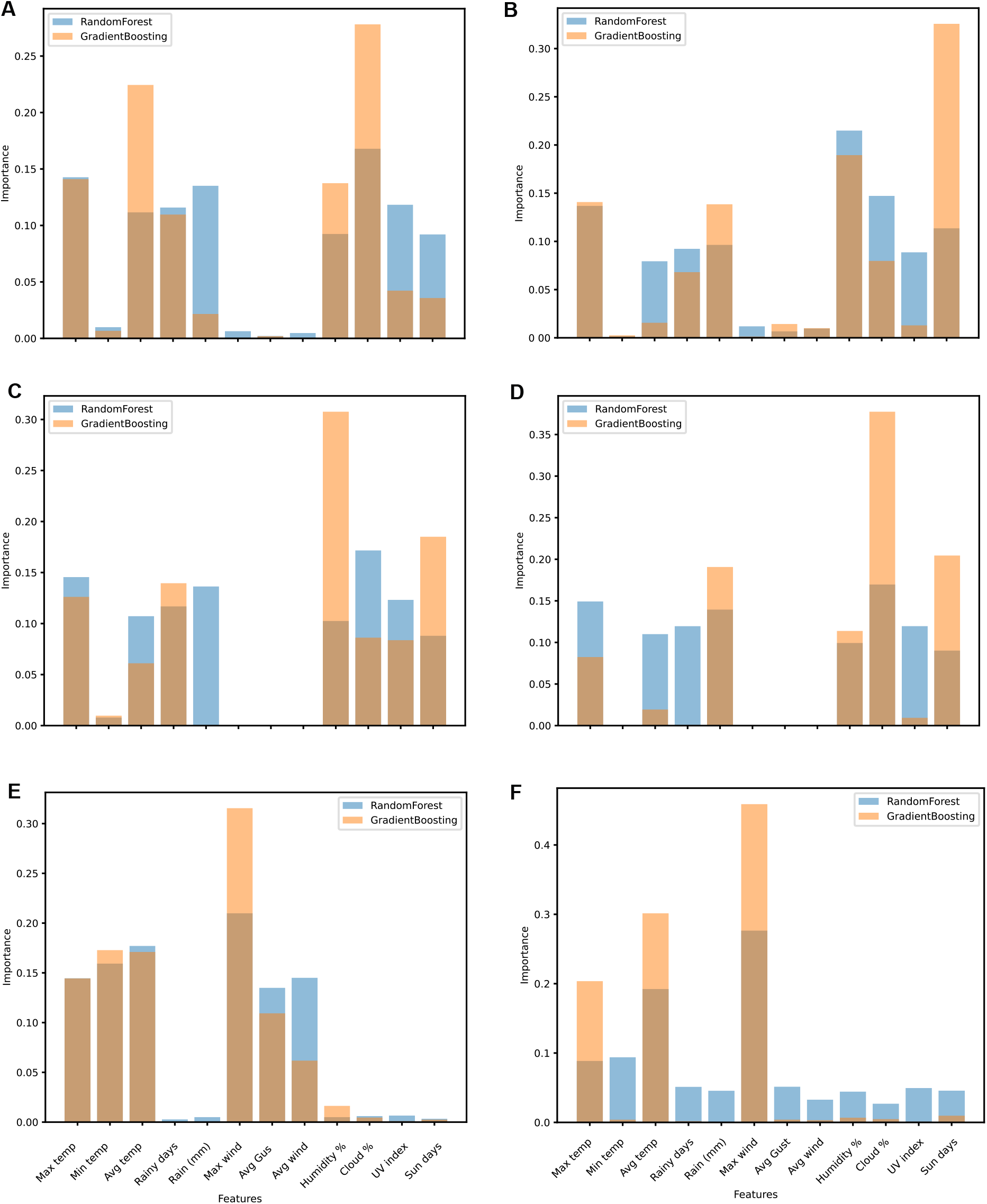
Trait-wise feature importance analysis. This barplot illustrates the trait-wise feature importances derived from Random Forest and Gradient Boosting models for predicting various plant traits. (A) - plant height; (B)-stem girth; (C) - inflo. branches; (D)-inflo. length; (E)-flowering date; (F) - plant maturity.

**Fig. S2:**
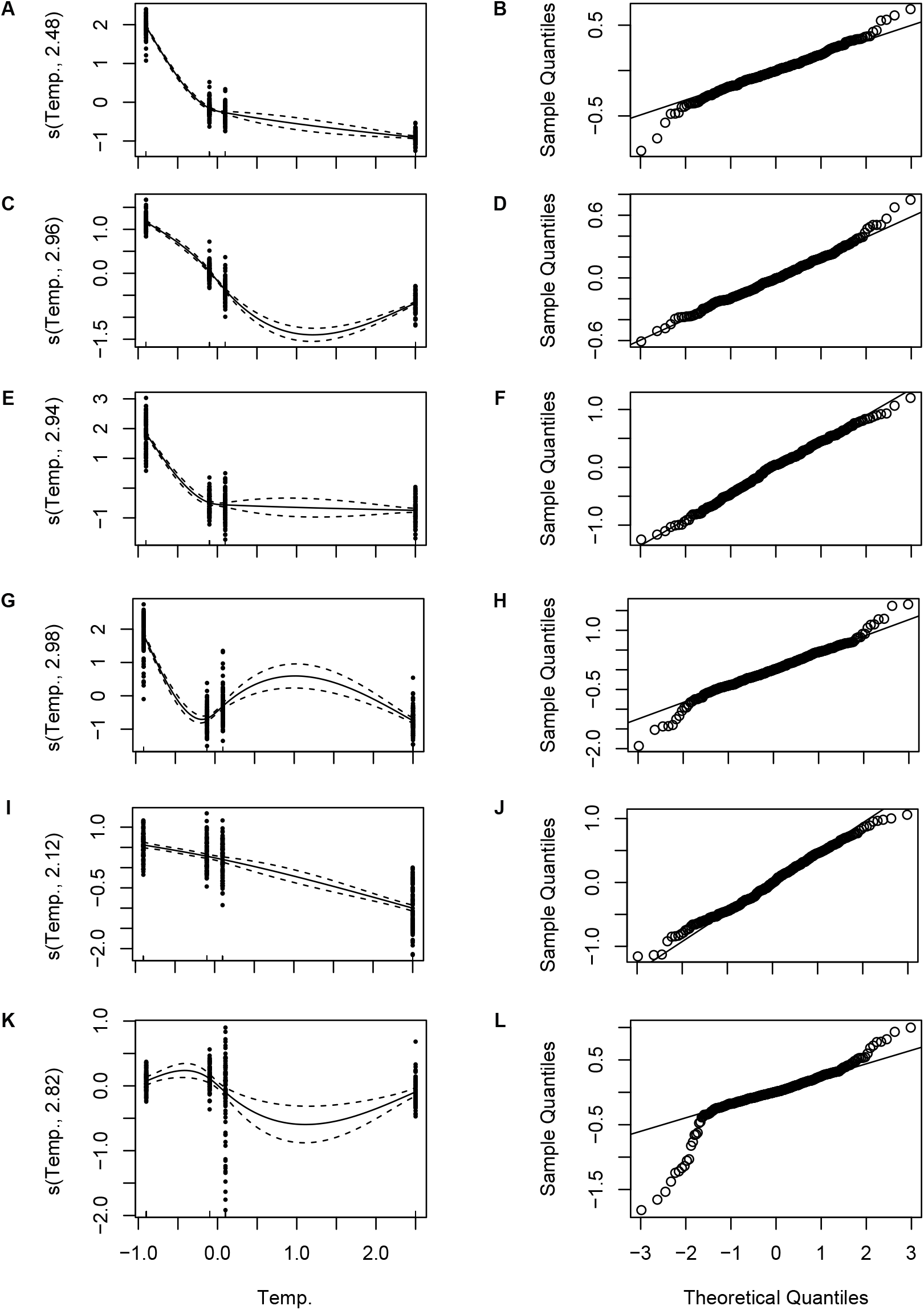
Residual and QQ plot analysis. Residual plot and QQ plot illustrating the distribution and normality of residuals from the fitted GAM, used to assess model adequacy and assumptions (A) & (B) - plant height; (C) & (D) - stem girth; (E) & (F) - inflo. branches; (G) & (H) - inflo. length; (I) & (J) - flowering date; (K) & (L) - plant maturity.

**Fig. S3:**
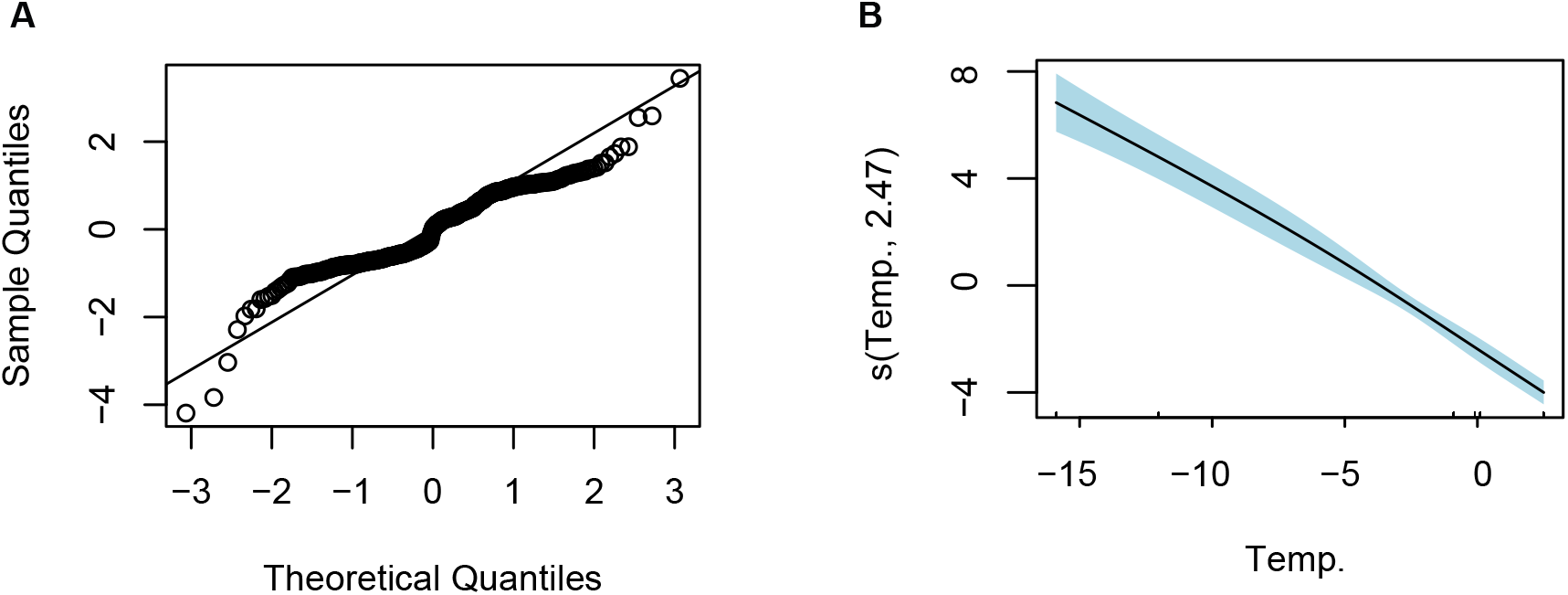
Residual normality and smooth effects analysis. (A) Quantile-Quantile (QQ) plot of the residuals from the transfer learning derived GAM model, illustrating the distribution of residuals compared to a theoretical normal distribution. (B) Smooth term plot showing the estimated relationship between the median-centered growth temperature and plant height. The shaded region represents the 95% confidence interval.

**Table S1:** List of *Amaranthus* genotypes.

|  |  |  |  |  |
| --- | --- | --- | --- | --- |
| Annapurna | IC-17930 | IC-38280 | IC-42415 | IC-95286 |
| DURGA | IC-17931 | IC-38282 | IC-42421 | IC-95299 |
| EC-146543 | IC-17932 | IC-38313 | IC-42776-2 | IC-95302 |
| EC-146546 | IC-17933 | IC-38353 | IC-42987-4 | IC-95304 |
| EC-17926 | IC-17935 | IC-38600 | IC-43715 | IC-95308 |
| EC-321563 | IC-17936 | IC-38611 | IC-467897 | IC-95313 |
| EC-359412 | IC-17937 | IC-38658 | IC-47436 | IC-95315 |
| EC-359417 | IC-17938 | IC-38661 | IC-82625 | IC-95316 |
| EC-519512 | IC-17947 | IC-38665 | IC-93946 | IC-95321 |
| EC-519523 | IC-21806 | IC-398214 | IC-94656 | IC-95322 |
| EC-519556 | IC-21938 | IC-415260 | IC-94661 | IC-95326 |
| IC-107127 | IC-21941 | IC-42258 | IC-95247 | IC-95330 |
| IC-107144 | IC-35407 | IC-42264 | IC-95249 | IC-95339 |
| IC-17923 | IC-35415 | IC-42346-6 | IC-95250 | PRA-2 |
| IC-17925 | IC-38131 | IC-42353 | IC-95251 |  |
| IC-17928 | IC-38185 | IC-42356 | IC-95253 |  |
| IC-17929 | IC-38193 | IC-42358 | IC-95277 |  |
|  | IC-38250 | IC-42397 | IC-95279 |  |
|  | IC-38269 | IC-42402 | IC-95284 |  |

**Table S2:** Generalized additive model (GAM) results.

|  | **Plant height** | **Stem girth** | **Inflorescence length** | **Inflorescence branches** | **Flowering date** | **Plant maturity** |
| --- | --- | --- | --- | --- | --- | --- |
| **Smooth term statistics** | edf = 2.978<br>Ref.df = 3<br>F-statistic = 4179<br>p-value <2e-16 | edf = 2.959<br>Ref.df = 3<br>F-statistic = 1637<br>p-value <2e-16 | edf = 2.981<br>Ref.df = 3<br>F-statistic = 768.1<br>p-value <2e-16 | edf = 2.94<br>Ref.df = 3<br>F-statistic = 916.9<br>p-value <2e-16 | edf = 2.115<br>Ref.df = 3<br>F-statistic = 265.4<br>p-value <2e-16 | edf = 2.823<br>Ref.df = 3<br>F-statistic = 7.239<br>p-value = 6.8e-05 |
| **Model fit statistics** | R-sq.(adj) = 0.958<br>Deviance explained = 96.9%<br>fREML = 27.99<br>est. = 0.071692 | R-sq.(adj) = 0.902<br>Deviance explained = 92.7%<br>fREML = 23.737<br>est. = 0.069941 | R-sq.(adj) = 0.784<br>Deviance explained = 84%<br>fREML = 253.24<br>est. = 0.39444 | R-sq.(adj) = 0.811<br>Deviance explained = 85.9%<br>fREML = 217.07<br>est. = 0.30384 | R-sq.(adj) = 0.778<br>Deviance explained = 83.4%<br>fREML = 227.69<br>est. = 0.34356 | R-sq.(adj) = 0.796<br>Deviance explained = 84.8%<br>fREML = 206.78<br>est. = 0.2848 |
| **AIC** | 48.75975 | 40.07758 | 648.9483 | 557.1366 | 599.7621 | 534.4562 |
| **BIC** | 404.212 | 395.5253 | 1004.401 | 912.5772 | 951.8575 | 889.7925 |

